# Condensin facilitates sister chromatid separation in actively transcribed DNA regions by relieving the obstructive effect of transcription

**DOI:** 10.1101/644799

**Authors:** Norihiko Nakazawa, Orie Arakawa, Mitsuhiro Yanagida

## Abstract

The evolutionarily conserved protein complex, condensin, is central to chromosome dynamics, including mitotic chromosome condensation and segregation. Genome-wide localization of condensin is correlated with transcriptional activity; however, the significance of condensin accumulation in transcribed regions remains unclear. Here, we demonstrate that condensin relieves the obstructive effect of mitotic transcription on sister chromatid separation in fission yeast, *Schizosaccharomyces pombe*. Time-lapse visualization of sister chromatid DNA separation revealed that mutant condensin causes delayed segregation specifically at mitotically transcribed, condensin-bound gene locus, *ecm33*^+^. Contrarily, the delay was abolished by transcriptional shut-off of the actively transcribed gene. We also showed that delayed separation at a heat shock-inducible gene locus, *ssa1*^+^, in condensin mutants was significantly alleviated by deletion of the gene. Since condensin has ability to remove ssDNA-binding proteins and RNA from unwound ssDNAs or DNA-RNA hybrids *in vitro*, we propose a model that condensin-mediated removal of mitotic transcripts from chromosomal DNA is the primary mechanism of sister chromatid separation.

## Introduction

In proliferative cells, mitotic chromosome condensation and subsequent segregation of replicated chromosomes into daughter cells are essential for faithful inheritance of genomic DNA. ‘Condensin’ has been named as an essential factor for mitotic chromosome condensation, originally demonstrated in mitotic frog egg extracts (Hirano and Mitchison, 1994; Hirano et al., 1997). Today, condensin is known as a key chromosome organizer for many types of chromosome regulation in interphase and mitosis (Hirano, 2016). In fission yeast, *Schizosaccharomyces pombe*, condensin temperature-sensitive (ts) mutants have been identified by detecting the failure of mitotic chromosome condensation in the same period of condensin discovery in the frog extracts(Hirano and Mitchison, 1994; Saka et al., 1994). Two condensin ts mutants, *cut3-477* and *cut14-208*, were originally isolated, and gene cloning identified two SMC (structural maintenance of chromosomes) family proteins, SMC4/Cut3 and SMC2/Cut14, as the responsible gene products (Hirano, 2006; Saka et al., 1994; Uhlmann, 2016). SMC subunits consist of two globular head domains at the N- and C-termini, which are connected by coiled-coil regions, interrupted by a central hinge region. The condensin Cut3-Cut14 dimer binds to three non-SMC condensin subunits (Cnd2/Barren and HEAT repeat-containing Cnd1 and Cnd3) forming the heteropentameric holo-complex (Sutani et al., 1999; Yoshimura et al., 2002).

Mapping of condensin binding sites at the whole chromosome level proposed a tight correlation between condensin association and transcription. Besides condensin enrichment at centromeres and RNA polymerase I (RNAP I)-transcribed ribosomal DNA (rDNA) repeats (Nakazawa et al., 2008; Wang et al., 2005), genomewide ChIP (Chromatin immunoprecipitation) analyses revealed that condensin associates with RNA polymerase III (RNAP III)-transcribed genes, such as tRNA genes, as well as retrotransposons in budding and fission yeasts (D’Ambrosio et al., 2008; Iwasaki et al., 2010; Tanaka et al., 2012). Fission yeast condensin also accumulates at genes highly transcribed by RNA polymerase II (RNAP II) in interphase and mitosis (Nakazawa et al., 2015; Sutani et al., 2015; Toselli-Mollereau et al., 2016). TATA box-binding protein, TBP, and mitotic transcription factors reportedly recruit condensin to RNAP II-transcribed, transcriptionally activated genes (Iwasaki et al., 2015; Kim et al., 2016). Thus, condensin binding at chromosomal DNA accompanies active transcription by RNAPs I, II, and III. Preferential binding to active genes has also been proposed for higher eukaryotic condensin I and II (Dowen et al., 2013; Kim et al., 2013; Kranz et al., 2013; Sutani et al., 2015), and even for prokaryotic condensin (Gruber and Errington, 2009), indicating that transcription-driven condensin accumulation is critical to understand the molecular action of condensin at mitotic chromosomes. However, physiological significance of condensin binding at actively transcribed DNA regions is still enigmatic.

In this study, we performed detailed, time-lapse, live cell analysis of sister chromatid DNA separation during fission yeast mitosis, and demonstrated that condensin ensures sister chromatid separation at mitotically upregulated gene and heat shock-inducible gene loci. Our results provide insight into how mitotic transcription is reconciled with chromosome segregation.

## Results and Discussion

### Sister chromatid DNA separation delays at mitotically transcribed gene locus in condensin mutant

We have previously shown that *S. pombe* condensin preferentially accumulates at mitotically transcribed genes and heat shock-inducible genes during mitosis (Nakazawa et al., 2015). To understand how condensin contributes to segregate transcriptionally activated regions in mitosis, we examined separation timing of the locus that binds to condensin. To visualise the locus, 7.8kb *lacO* arrays were chromosomally integrated within 2 kb of the mitotically up-regulated *ecm33*^+^ gene locus (Fig. 1A, left) in wild-type and temperature-sensitive (ts) *cut14-208* mutant cells. LacI-GFP was expressed in these cells, and *lacO* arrays that bound to LacI-GFP were visualized (Nabeshima et al., 1998). *lacO* arrays were also integrated along the long arm of chromosome 1, within 2 kb of four other genes (*sod2*^+^, SPAC4G8.03c, *sck1*^+^, and *nsk1*^+^), which were not transcriptionally activated during mitosis. The level of bound condensin was negligible at the four genes (Nakazawa et al., 2015). To visualise the spindle pole body (SPB; equivalent to a centrosome), we used Sad1-mCherry, which is located at the SPBs (Fig. 1A, right) (Hagan and Yanagida, 1995). To monitor the time-course of *lacO* segregation, cultures grown at 26°C were shifted to 36°C (the restrictive temperature for the *cut14-208* mutation in the coiled-coil region, S861P) (Sutani and Yanagida, 1997), and >40 mitotic cells were analyzed by taking movies for each *lacO*/LacI-GFP locus in individual wild-type and *cut14-208* cells.

**Figure 1.**
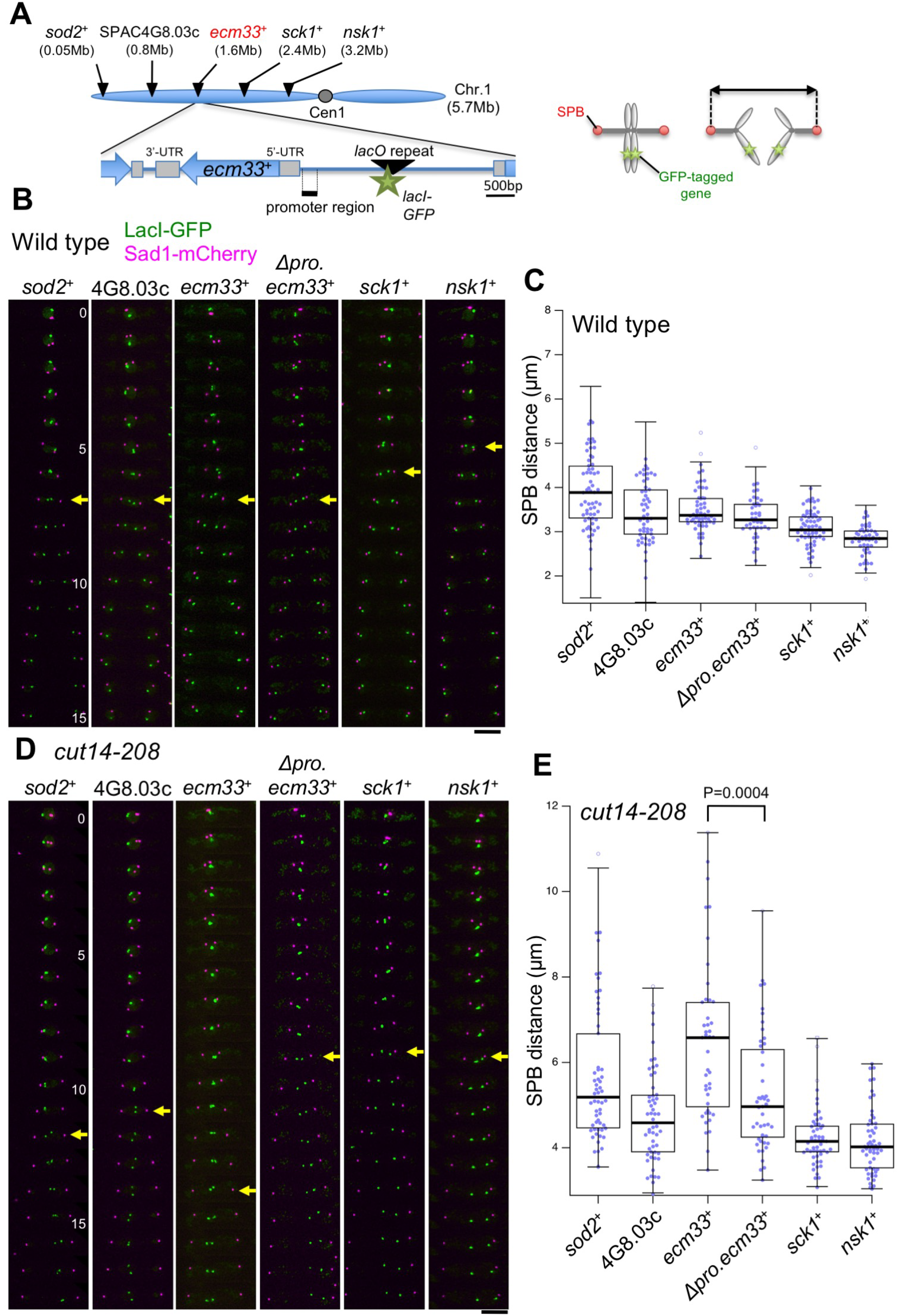
Delay in sister locus separation at mitotically transcribed *ecm33*^+^ in condensin mutant, and restoration of the delay by removing the *ecm33*^+^ promoter. **(A)** (Left) Schematic diagrams of the *ecm33*^+^ locus and positions of inserted *lacO* on chromosome 1. Black triangles indicate five distinct positions of the *lacO* arrays. The distance (in Mb) from the left telomere is shown in parentheses under each locus. In the *ecm33* promoter-less mutant (*Δpro ecm33*^+^), a 0.2 kb sequence of the 5’-UTR was replaced with the hygromycin resistance gene. (Right) Scheme for measurement of the distance between two SPBs (bi-directed arrow) that determined the timing of sister locus separation. **(B, D)** Time-lapse frames of projected images were obtained using a Delta Vision microscope. Loci (*lacO*/LacI-GFP, green) and SPBs (Sad1-mCherry, magenta) were observed in living wild-type (B) and *cut14-208* (C) mutant cells cultured at 36°C for 1.5 h. Selected frames are shown every 1min. Yellow arrowheads indicate timing of sister GFP locus separation (when signals were separated >0.5 µm). Scale bar, 5 µm. **(C, E)** Box plots of measured distances between SPBs when sister GFP locus signals were separated. At each locus, movies of more than 40 mitotic cells were analyzed. The center line indicates the median, and the box limits are from the second to the third quartile (25% to 75% of the data points). The whiskers extend from the box limits to the 2nd and 98th percentile values, with outliers and far outliers indicated by white circles and squares outside the whiskers, respectively. The *P* value from a two-tailed Wilcoxon-Mann-Whitney test is shown. Statistical data are provided in Supplemental Table 1. Transcription of *ecm33*^+^ gene caused a delay in sister locus separation during mitosis in the condensin mutant.

Sister GFP loci (green dots) were separated smoothly in wild-type cells, when the SPB distance (between magenta SPB dots) reached 3-4 µm (Fig. 1B, arrows). Sister loci were considered separated when GFP signals were fully split (> 0.5 µm). SPB distances were then determined at the time point in which the two GFP-dots separated. Measurements from a number of movies (~50 cells at each locus, total 635 movies, statistical data is shown in Supplemental Table 1) revealed an inverse relationship between the distance of the loci from the telomere and the mean SPB distance for GFP signal splitting (median bars in Fig. 1C). The telomere-proximal *sod2*^+^ locus took longer to split than the centromere-proximal *nsk1*^+^ locus.

Representative time-lapse images for *cut14-208* condensin mutant cells are shown in Fig. 1D. The SPB distance at the time of sister-locus separation for *sod2*^+^, SPAC4G8.03c, *sck1*^+^, and *nsk1*^+^ was slightly longer (4-5 µm) than in WT (3-4 µm). On the other hand, the SPB distance for splitting of the mitotically transcribed *ecm33*^+^ locus was strikingly longer. Splitting did not occur until the SPB distance reached a mean of 6.5 µm in mutant cells. Thus, the mitotically highly transcribed *ecm33*^+^ gene locus showed an anomalous segregation delay in condensin mutant cells, relative to the 4 control loci (Fig. 1E).

### Transcriptional shut-off abolishes the delay of sister chromatid DNA separation at actively transcribed genes in condensin mutant

To examine whether transcription at *ecm33*^+^ affects separation timing at this locus, a promoter-less mutant, *Δpro ecm33*, was constructed. *ecm33*^+^ expression was greatly diminished (~20%) in *Δpro ecm33* compared to that in WT cells, as measured by reverse transcription-PCR (Supplemental. Fig. S1). In *Δpro ecm33* cells, the delay of separation at the *ecm33*^+^ locus was essentially abolished (the mean SPB distance became ~5µm; Fig. 1E), suggesting an opposing effect of mitotic transcription on sister chromatid DNA separation.

We have also visualized an hsp gene locus, *ssa1*^+^, by integrating *lacO* arrays 6 kb upstream of the gene (Fig. 2A). In WT cells, separation timing of the *ssa1*^+^-proximal *lacO* DNAs was not significantly changed in the deletion mutant of *ssa1*^+^ (*Δssa1*) after heat shock at 36°C (Fig. 2B, Supplemental Table 2). Contrarily, separation timing at that locus became significantly earlier by deleting the *ssa1*^+^ gene in *cut14-208* (Fig. 2C). We thus concluded that in the condensin mutant, transcription *per se* caused the delay in sister chromatid DNA splitting at both mitotically-activated and heat-shock responsible gene loci during mitosis.

**Figure 2.**
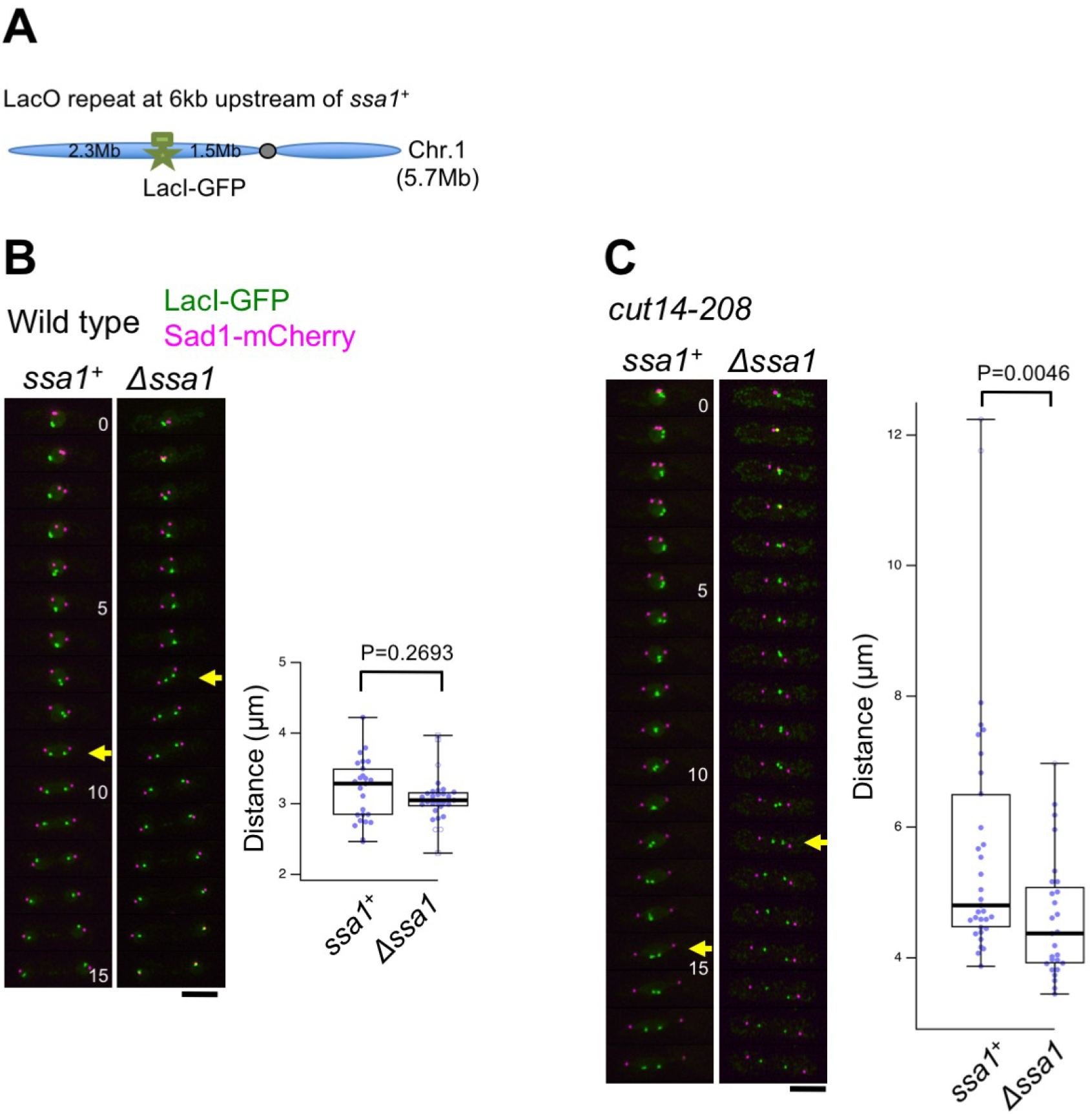
Deletion of a heat shock-inducible *ssa1*^+^ gene alleviated the delay of sister DNA splitting at the locus in *cut14-208* mutant. **(A)** A diagram of the *ssa1*^+^ locus and the position of the inserted *lacO* on chromosome 1. For constructing the *Δssa1* mutant, the ORF sequence of *ssa1*^+^ was replaced with the hygromycin resistance gene. **(B, C)** (Left) Time-lapse frames of projected images were obtained using a Delta Vision microscope. The gene loci (*lacO*/LacI-GFP, green) and the SPBs (Sad1-mCherry, magenta) were observed in living wild-type (B) and *cut14-208* (C) mutant cells cultured at 36°C for 1.5 h. Images taken at 1 min intervals are shown. Yellow arrowheads indicate the timing of sister GFP locus separation (when signals were separated >0.5 µm). Scale bar, 5 µm. (Right) Box plots of measured distances between SPBs in wild-type (B) and *cut14-208* (C) mutant cells. At each locus, movies of more than 25 mitotic cells were analyzed. The center line indicates the median, and the box limits are from the second to the third quartile (25% to 75% of the data points). The whiskers extend from the box limits to the 2nd and 98th percentile values, with outliers and far outliers indicated by white circles and squares outside the whiskers, respectively. *P* values from a two-tailed Wilcoxon-Mann-Whitney test are shown. Statistical data are provided in Supplemental Table 2.

### Condensin-mediated elimination of RNA transcripts may facilitate mitotic chromosome segregation

The results indicate that condensin negates the inhibitory effect of mitotic transcription on chromosome segregation at actively transcribed gene loci. If we consider *in vitro* activity of the condensin SMC heterodimer (Cut3-Cut14), it promotes the DNA renaturation reaction which winds up of complementary ssDNA to dsDNA (Sutani and Yanagida, 1997). In addition, a ssDNA-binding protein, Ssb1/RPA, bound to ssDNA, and also RNA molecules hybridized to ssDNA were removed by condensin SMC through the DNA renaturation reaction (Akai et al., 2011). This activity is strikingly reduced in condensin ts mutants, *cut14-208* (coiled-coil mutation, S861P) or *cut14-y1* (hinge mutation, L543S), implying that RNA transcripts bound at actively transcribed regions interfere with chromosome segregation. Transcriptional attenuation using transcription inhibitors or mutation in the transcription mediator protein suppresses chromosome segregation defects in condensin mutants (Supplemental Fig. S2) (Sutani et al., 2015). Taken together, we now propose that condensin-mediated elimination of RNA transcripts on transcribed DNA regions facilitates mitotic chromosome segregation. Actually, many genes are actively transcribed even during mitosis. Our results provide a novel insight into mitotic chromosome segregation, which may be accomplished by removing obstructive RNA and/or proteins on chromosomes, in addition to forming condensed mitotic chromosomes. Further analysis is definitively required to understand the inhibitory effect of bound RNA on chromosome segregation during mitosis.

Previously, we found that *S. pombe* condensin enriches at central centromeric DNAs where kinetochores are assembled with specialized nucleosomes containing a centromere-specific histone H3 variant, CENP-A/Cnp1 (Allshire and Ekwall, 2015; Nakazawa et al., 2008; Takahashi et al., 2000). In addition to the pericentromeric outer repeat regions (otr), the central centromere also generates a non-coding RNA transcribed by RNAP II (Chan and Wong, 2012; Choi et al., 2011; Talbert and Henikoff, 2018). Condensin enrichment at the central centromere appears to be dependent on transcription (Supplemental Fig. S3), implying that condensin acts on centromeric transcripts. In metaphase, replicated sister centromeric DNAs are pre-separated for establishment of bi-oriented kinetochore-spindle attachment (Nabeshima et al., 1998); however, the pre-separation fails in condensin mutant cells, resulting in unequal segregation of centromere-proximal sister DNAs (Nakazawa et al., 2008). Condensin may play a crucial role in separating centromeres by removing non-coding RNA transcripts, as well as in actively-transcribed chromosomal arm regions.

## Materials and Methods

### Strains, plasmids, and media

The *S. pombe* haploid wild-type strain 972 *h*^−^ and its derivative mutant strains, including the temperature-sensitive (ts) *cut14-208*, *cut3-477* and *cnd2-1* were used (Aono et al., 2002; Saka et al., 1994). Disruption of the *ecm33*^+^ promoter (Takada et al., 2010) was carried out using a two-step PCR-based gene targeting method (Bähler et al., 1998). The *ecm33*^+^ gene was cloned into a pSK248 plasmid under the native promoter and expressed in an *ecm33*^+^ promoter-less mutant to alleviate the cell morphology defect (Takada et al., 2010).

Labelling of chromosomal loci at *ecm33*^+^, *sod2*^+^, SPAC4G8.03c, *sck1*^+^, *nsk1*^+^, and *ssa1*^+^ using the lac repressor (LacI)/ lac operator (*lacO*) system was performed as previously described (Nabeshima et al., 1998), with the following modifications. The pMK19A-KanR plasmid derived from pMK2A was used for *lacO* insertion at each locus. pMK19A-KanR contains the *lacO* repeat (7.8 kb) and the kanamycin-resistance gene *Kan*^*R+*^ (1.7 kb fragment from pFA6a-KanMX) (Bähler et al., 1998). Target sequences (1.5-2.0 kb) were then cloned into pMK19A-KanR. Resulting plasmids were linearized by cleavage within the target sequences and transformed into a strain expressing the GFP-LacI-NLS protein.

Thiolutin (Sigma T3450) and 1,10-phenanthroline (Sigma 320056) were diluted with dimethyl sulphoxide (1 mg/mL) and ethanol (30 mg/mL), respectively, and used to inhibit RNA polymerases I, II, and III (Lackner et al., 2007). Culture media used for *S. pombe* were YPD and SPA sporulation medium (Saka et al., 1994). Cells were counted using a haematology analyzer (Sysmex FDA-500).

### Live cell analysis

*S. pombe* cells were cultured at 26°C in EMM2 medium and shifted to 36°C for 1.5 hr. Before observation, exponentially growing cells were transferred to a glass-bottomed dish (IWAKI Glass) coated with 10 mg/mL concanavalin A (Wako). Time-lapse images were recorded with a DeltaVision microscope system (Applied Precision, GE Healthcare UK Ltd). The objective lens was an oil immersion lens (PlanApo 60x, NA 1.4; Olympus). For observations of *lacO*/LacI-GFP and Sad1-mCherry signals, a set of images from 7 focal planes at 0.2-µm intervals was obtained every 0.5 min. Image projection and deconvolution were performed using an imaging workstation (SoftWoRx; Applied Precision). More than 40 mitotic cells (for Figure. 1) or 25 (for Figure. 2) were observed for each locus. Image J (NIH) was used to measure the SPB distance when *lacO*/LacI-GFP signals were separated. We defined the separation timing of sister *lacO*/LacI-GFP signals when the distance between proximal edges of the GFP signals was greater than 0.5 µm.

### RNA extraction, reverse transcription PCR, and chromatin immunoprecipitation

Total RNA from *S. pombe* cells was extracted using the hot-phenol method. One µg of total RNA was reverse-transcribed using PrimeScript RT reagent (TaKaRa) with oligo dT primers. A genomic DNA eraser supplied with reverse transcription reagent was used to remove contaminating genomic DNA in the RNA sample. cDNA solution was diluted 25-fold with RNase-free water (Ambion) and 5 µL were used for PCR. Results were quantified using real-time PCR with SYBR premix Ex Taq II solution (TaKaRa). Chromatin immunoprecipitation (ChIP) was performed as previously described (Nakazawa et al., 2015).

## Acknowledgments

We thank Dr. Steven D. Aird for editing the manuscript. NN was supported by a Grant-in-Aid for Young Scientists (B) 25840013 from the Japan Society for the Promotion of Science (JSPS). We are also grateful for the generous support of OIST.

---

**Supplemental Figure S1.**
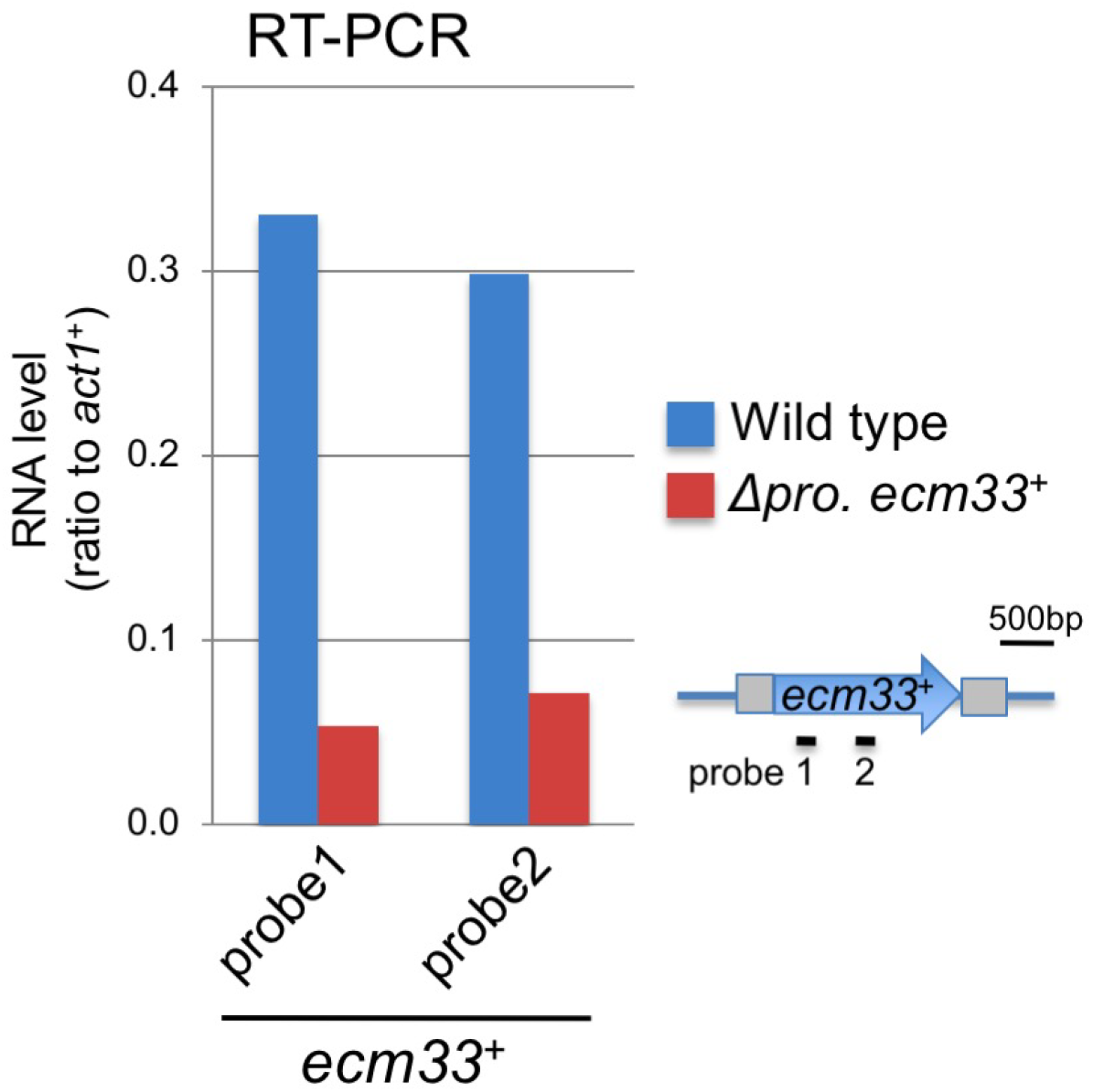
The *ecm33*^+^ gene transcript level was reduced in the promoter-less mutant, *Δpro ecm33*^+^. Transcript levels of *ecm33*^+^ were determined in asynchronously cultured WT and promoter-less mutant *Δpro ecm33*^+^ cells using quantitative reverse-transcription (RT)-PCR. Results were obtained using two probes (probes 1 and 2).

**Supplemental Figure S2.**
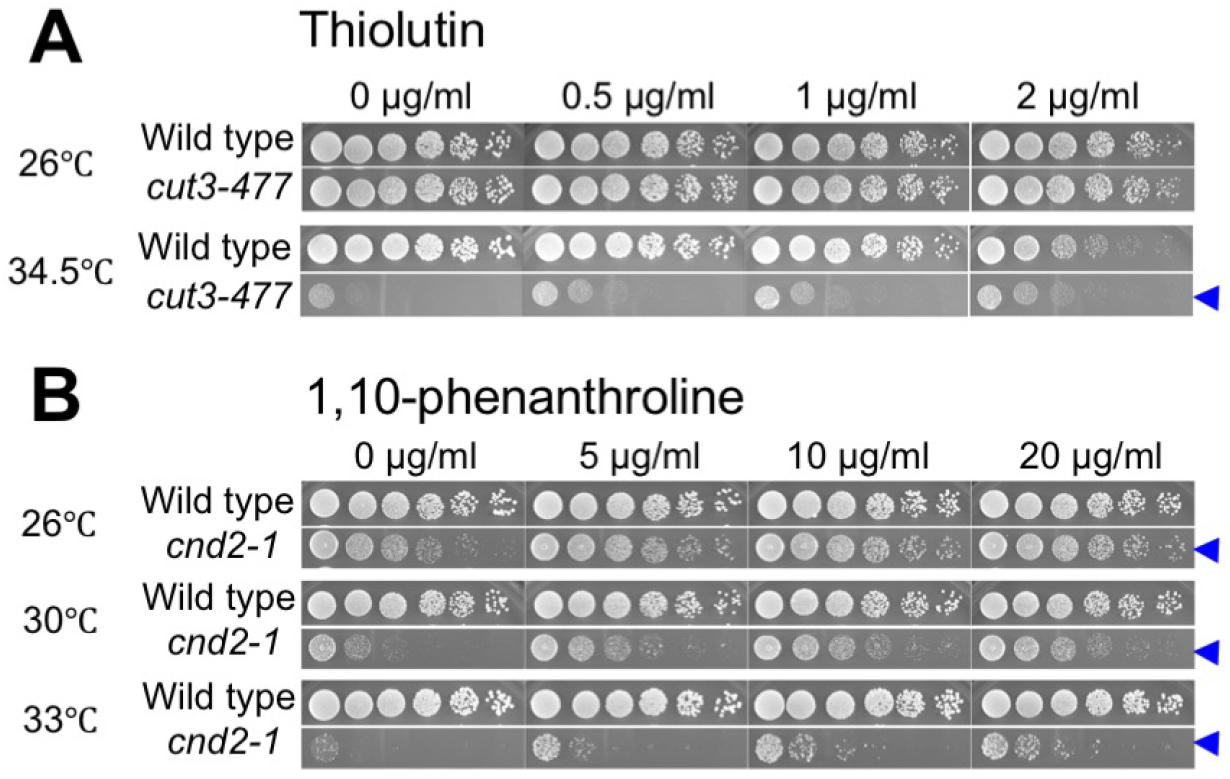
Partial rescue of temperature-sensitive condensin mutants by transcription inhibitors. The transcription inhibitors thiolutin and 1,10-phenanthroline partly suppressed the temperature-sensitive (ts) phenotype of *cut3-477* and *cnd2-1* mutants at 34.5°C and 26-33°C, respectively (arrowheads). Both drugs target RNA polymerases I, II, and III. These data confirm the results of Sutani and colleagues (Sutani et al., 2015).

**Supplementary Figure S3.**
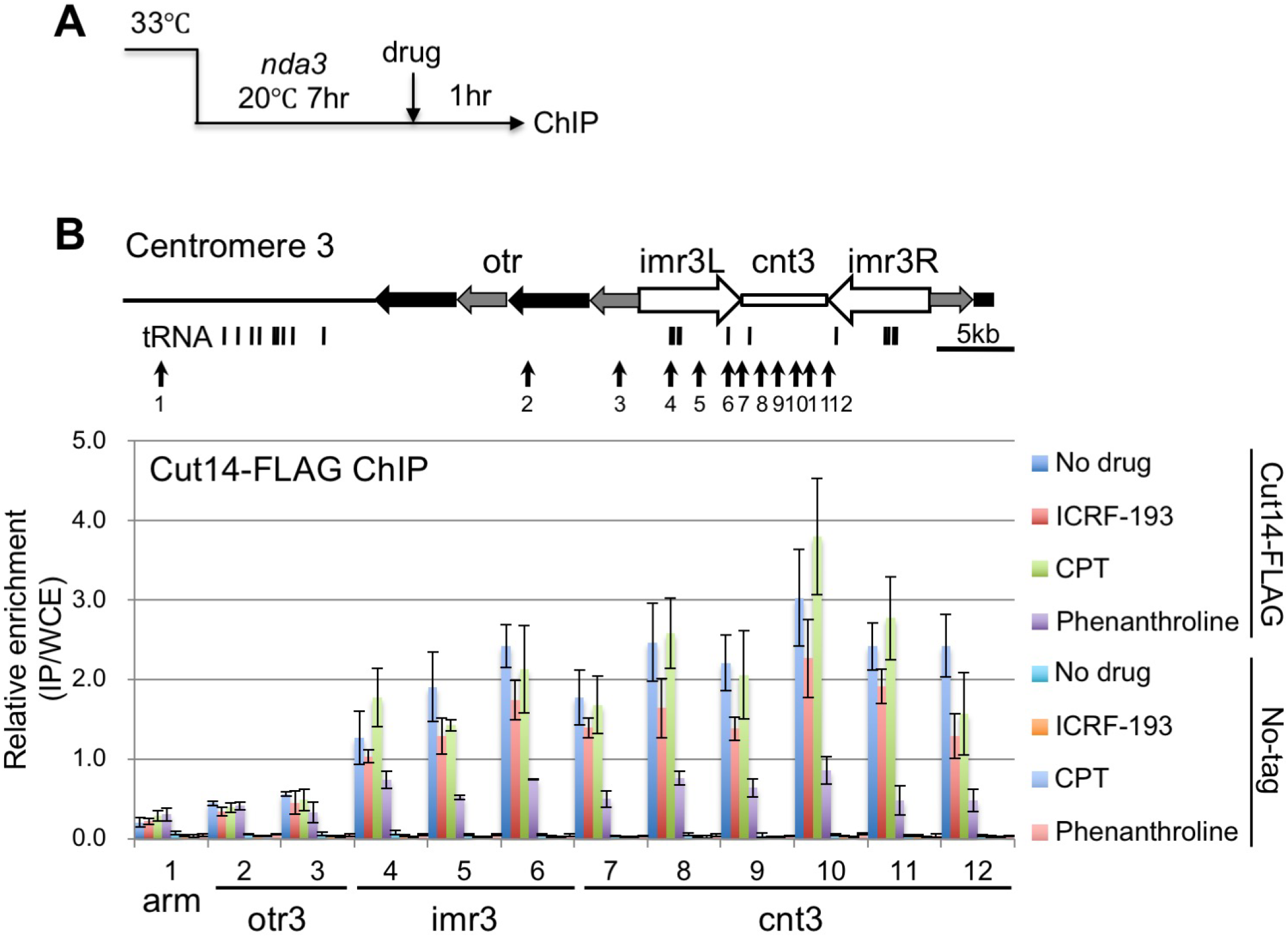
Centromeric association of condensin decreased in the presence of a transcription inhibitor. ChIP (Chromatin immunoprecipitation)-quantitative PCR analysis of Cut14-FLAG was performed at the centromeric region of chromosome 3 in *nda3-KM311* β-tubulin cs mutants (Hiraoka et al., 1984), along with a non-tagged control strain. **(A)** Procedure of experiment. Cells were mitotically arrested at 20°C for 7 hr, and then incubated at 20°C for additional 1hr in the presence of 50 µM ICRF-193 (DNA topoisomerase II inhibitor) (Nakazawa et al., 2016), 50 µM CPT (Camptothecin, DNA topoisomerase I inhibitor), or 120 µg/ml 1,10-phenanthroline (Transcription inhibitor). DMSO was used for a mock treatment. **(B)** ChIP results. Positions of PCR primers (arrows with numbers) are indicated in a schematic illustration of centromere 3. Relative enrichment was calculated as the ratio of IP to WCE with error bars showing standard deviations (n=3). Association of Cut14 at central centromeric DNA (cnt3 and imr3) was remarkably decreased by transcriptional inhibition, and partly affected by inhibition of DNA topoisomerase II.

**Supplemental Table 1.** Statistical data of measured distances between SPBs when sister GFP locus signals separated (*ecm33*^+^, *Δpro ecm33*^*+*^, *sod2*^+^, SPAC4G8.03c, *sck1*^+^, and *nsk1*^+^ loci).

| Strain | Wild type |  |  |  |  |  | *cut14-208* |  |  |  |  |  |
| --- | --- | --- | --- | --- | --- | --- | --- | --- | --- | --- | --- | --- |
| Locus | *sod2*<sup>+</sup> | SPAC<br>4G8.03c | *ecm33*<sup>+</sup> | *Δpro.*<br>*ecm33*<sup>+</sup> | *sck1*<sup>+</sup> | *nsk1*<sup>+</sup> | *sod2*<sup>+</sup> | SPAC<br>4G8.03c | *ecm33*<sup>+</sup> | *Δpro.*<br>*ecm33*<sup>+</sup> | *sck1*<sup>+</sup> | *nsk1*<sup>+</sup> |
| Number of cells | 61 | 53 | 56 | 40 | 67 | 48 | 58 | 54 | 43 | 47 | 53 | 55 |
| Minimum | 2.16 | 1.96 | 2.44 | 2.34 | 2.02 | 1.93 | 3.55 | 2.91 | 3.48 | 3.24 | 3.07 | 3.03 |
| 25th percentile | 3.30 | 2.94 | 3.22 | 3.07 | 2.88 | 2.66 | 4.48 | 3.90 | 4.95 | 4.24 | 3.89 | 3.52 |
| Median | 3.89 | 3.30 | 3.37 | 3.27 | 3.04 | 2.81 | 5.19 | 4.59 | 6.58 | 4.97 | 4.15 | 4.02 |
| 75th percentile | 4.50 | 3.96 | 3.76 | 3.63 | 3.35 | 3.03 | 6.48 | 5.24 | 7.41 | 6.31 | 4.52 | 4.57 |
| Maximum | 5.51 | 4.65 | 5.24 | 4.90 | 3.98 | 3.45 | 10.88 | 7.78 | 11.38 | 9.55 | 6.58 | 5.97 |
| Average | 3.92 | 3.44 | 3.52 | 3.33 | 3.09 | 2.81 | 5.65 | 4.75 | 6.49 | 5.26 | 4.24 | 4.18 |
| S.D. | 0.79 | 0.65 | 0.51 | 0.49 | 0.38 | 0.34 | 1.60 | 1.12 | 1.87 | 1.40 | 0.68 | 0.80 |

Time-lapse images were recorded by the DeltaVision microscope system (Applied Precision, GE Healthcare UK Ltd). Image projection and deconvolution were performed using an imaging workstation (SoftWoRx; Applied Precision). Image J (NIH) was used to measure the SPB distance when *lacO*/*LacI*-GFP signals were separated greater than 0.5 μm (Materials and methods). The unit is μm except for number of cells.

**Supplemental Table 2.** Statistical data of measured distances between SPBs when sister GFP locus signals separated (*ssa1*^+^ and *Δssa1*^+^ loci).

| Strain | Wild type |  | *cut14-208* |  |
| --- | --- | --- | --- | --- |
| | *ssa1*<sup>+</sup> | Δ*ssa1* | *ssa1*<sup>+</sup> | Δ*ssa1* |
| Number of cells | 25 | 29 | 33 | 27 |
| Minimum | 2.47 | 2.30 | 3.87 | 3.45 |
| 25th percentile | 2.84 | 2.96 | 4.47 | 3.92 |
| Median | 3.29 | 3.05 | 4.80 | 4.37 |
| 75th percentile | 3.50 | 3.16 | 6.51 | 5.09 |
| Maximum | 4.22 | 3.97 | 12.24 | 6.97 |
| Average | 3.19 | 3.07 | 5.71 | 4.59 |
| S.D. | 0.44 | 0.33 | 2.00 | 0.93 |

Time-lapse images were recorded by the DeltaVision microscope system (Applied Precision, GE Healthcare UK Ltd). Image projection and deconvolution were performed using an imaging workstation (SoftWoRx; Applied Precision). Image J (NIH) was used to measure the SPB distance when *lacO*/LacI-GFP signals were separated greater than 0.5 μm (Materials and methods). The unit is μm except for number of cells.

